# Cross-species transmission of human hepatitis B virus to wild Neotropical primates

**DOI:** 10.1101/2024.10.31.621242

**Authors:** Jean P. Boubli, Hani R. El Bizri, Luan F. Botelho-Souza, Chrysoula Gubili, Stephen J. Martin, Maisa da S. Araújo, Tommy C. Burch, Mariluce R. Messias, Alcione de O. dos Santos, Luiz S. Ozaki, André V.C. Pereira, Tony H. Katsuragawa, Ana Maísa Passos-Silva, Luiz H. S. Gil, Izeni P. Farias, Juan M.V. Salcedo, Deusilene Vieira

## Abstract

Hepatitis B virus (HBV) infects approximately one-third of the world’s human population and kills over one million people each year. HBV is also prevalent in Old World apes but not in New World primates. Human-to-primate transmission of HBV was suspected in zoo-captive monkeys and Mauritius macaques, but empirical data are scarce. Here, we collected blood and liver samples from 88 monkeys of 27 species in two areas of the Amazon, one with pristine forest and the other highly occupied and deforested by humans. A total of 17 (34.7%) out of 49 specimens from the human-occupied region tested positive for HBV. At this site, there was a positive relationship between human population density in the sampling location and the likelihood of primates being infected by HBV. Conversely, all 39 samples from the pristine forest tested negative for HBV. By sequencing a portion of the HBV S gene in five positive samples, each from a distinct primate genus, we found that four samples were closely related to the globally widespread human HBV-A strain, but not to the Americas-native HBV-F strain. The fifth sample aligned with the human HBV-D type, prevalent in the region where these samples were obtained. To our knowledge, this study represents the first reported cases of HBV in multiple wild New World primate species anywhere in the world. Our results suggest that primates were infected by strains brought into this part of Brazil by human immigrants, where HBV transmission may have been facilitated by the close contact between humans and monkeys due to high human occupation. This shows that the impact of human immigration, occupation and population growth in the Amazon extends beyond habitat loss; it also facilitates cross-species infections, potentially leading to the emergence of new, virulent viral strains that threaten both Amazonian biodiversity and human health.

**Author Summary:** Jean P. Boubli: Professor in Primate Ecology and Evolution, School of Science, Engineering and the Environment, University of Salford, Uk

Hani R. El Bizri: PhD in Wildlife Conservation with a focus on Amazonian sustainable development. Analyst of Center for International Forestry Research (CIFOR), Bogor, Indonesia

Luan F. Botelho-Souza: Master’s and doctorate in experimental biology at the Universidade Federal de Rondônia/Brazil,

Chrysoula Gubili: Researcher at the Fisheries Research Institute Nea Peramos, Kavala, Greece, specializing in population ecology and conservation genetics.

Stephen J. Martin: Professor in Social Insects, School of Science, Engineering and the Environment, University of Salford, Uk

Maisa da S. Araújo: Master’s and doctorate in experimental biology, at the Universidade Federal de Rondônia/Brazil,

Mariluce R. Messias: Professor of Zoology and curator of the Mammalogy Museum at the Universidade Federal de Rondônia/Brazil.

Alcione de O. dos Santos: Master’s and doctorate in experimental biology at the Universidade Federal de Rondônia/Brazil.

Luiz S. Ozaki: Master’s degree from Universidade de Brasília and PhD in Physiological Sciences, Molecular Biology in Fukuoka, Kyushu, Japan.

André V.C. Pereira: Bachelor in Biology the Universidade Federal de Rondônia/Brazil.

Tony H. Katsuragawa: Master’s and doctorate in experimental biology, at the Universidade Federal de Rondônia/Brazil.

Ana Maísa Passos-Silva: Master’s degree in experimental biology from the Federal University of Rondônia/Brazil.

Luiz H. S. Gil: Master’s in experimental biology, at the Universidade Federal de Rondônia/Brazil.

Izeni P. Farias: Professor of Genetics, Universidade Federal do Amazonas, Brazil

Juan M.V. Salcedo: Master in Tropical Medicine from the University of Brasilia/Brazil and PhD in Sciences from the University of São Paulo/Brazil.

Tommy C. Burch: Masters in Biological Sciences and PhD candidate from the University of Salford, UK.

Deusilene Vieira: Master in Tropical Medicine from the University of Brasilia/Brazil and PhD in Sciences from the University of São Paulo/Brazil.

## Introduction

Globalization has significantly accelerated the emergence and spread of infectious diseases, viral pathogens in particular [1]. Key examples include HIV, SARS, SARS-CoV-2, and Ebola, all of which have allegedly undergone spillovers from animals to humans. These cross-species transmissions, facilitated by increased global travel, trade, and environmental changes, have led to widespread health crises and substantial economic repercussions.

Our long-term close associations with animals also allow viral diseases to move in the other direction (from humans to animals), although very few cases are known except for polio [2], and most involve human to ape transmission [3]. Currently, two billion people are infected with Hepatitis B virus (HBV), which causes almost one million deaths each year [4]. It has been proposed that HBV first appeared in humans around 40,000 years ago via spillover from an unknown non-primate source (e.g., a rodent or bird), before jumping from humans to apes via independent transmission events involving chimpanzees, gibbons, and orangutans [5]. Human HBV is currently classified into eight distinct genotypes (A to H) [6], and the Old-World apes (gibbons, orangutans, chimpanzees, and gorillas) carry a group of distinctive nonhuman HBV genotypes [7-12]. Another HBV strain grouped with human genotype D, and it was isolated from free ranging long tail macaques (*Macaca fascicularis*) which were introduced into Mauritius [13]. However, according to previous extensive surveys [14] there have only been three reported cases of HBV in wild or captive New World primates (or monkeys) [7, 13]. One case was caused by a highly divergent form of HBV (WMHBV) found in a captive woolly monkey, *Lagothrix lagotricha,* at Louisville Zoo, USA, which is phylogenetically basal to all primates and intermediate between rodent and primate genotypes [15]. The other HBV strain grouped with human genotype D and it was isolated from free ranging long tail macaques (*M. fascicularis*), which were introduced into Mauritius [13].

More recently, a novel and highly divergent HBV species (CMHBV) was discovered in Brazilian capuchin monkeys (*Sapajus xanthosternos*) [16], which shares key functional similarities with human HBV, shedding light on the potential long-term coevolution of hepadnaviruses in primates.

New World primates have been in the Americas since the Oligocene [17], but humans only arrived on this continent ca 20,000 years ago [5] and they brought the HBV-F genotype, which remains restricted to the Americas [5]. A recent survey of HBV genotypes in the southwestern Amazon region of Brazil, an area that has undergone large-scale deforestation, indicated that HBV-A and -D genotypes are dominant due to the immigration of Africans and Europeans [18]. In this area, people have very close relationships with wild monkeys via wild meat hunting and the pet trade. Hence, it is possible that HBV may have crossed over to wild nonhuman primates in regions where human occupation of the Amazon forest was occurring. We tested this hypothesis by searching for HBV in a wide range of Amazonian primates from 11 species in two distinct sites of the Amazon region, one with very pristine forests, and one with high levels of human occupation and deforestation.

## Results

None of the 39 liver tissues from 11 species of monkeys collected from the pristine upper Japura River site tested positive for HBV. However, 17 (34.7%) blood samples from primates collected in the highly occupied and deforested site in southwestern Amazonia tested positive for HBV. HBV has been detected in primates at most of the capture sites (Fig 1), but our GLMM showed that the likelihood of a primate being positive for HBV was higher for capture sites with higher human population density (Coefficient: 0.13; t-value: 2.11; p-value: 0.039).

**Fig 1.**
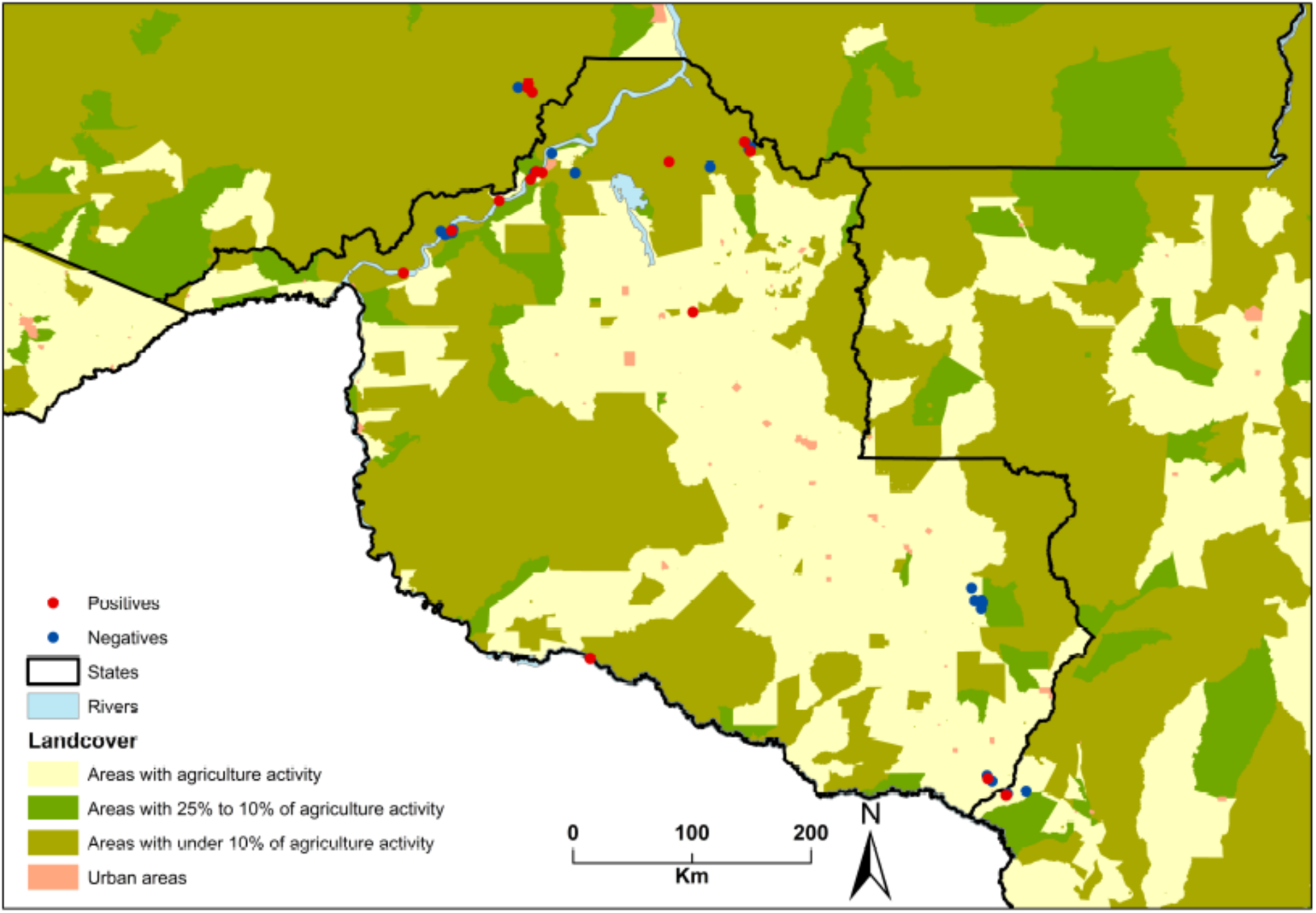
Map of Rondônia State in Brazil showing degree of land cover and the sampling locations, from which HBV was present or absent in the samples.

From the 17 positive samples, five were chosen for sequence analysis that represented primates from five different genera. The presence of HBV was reconfirmed before the products were sequenced (Fig 2).

**Fig 2.**
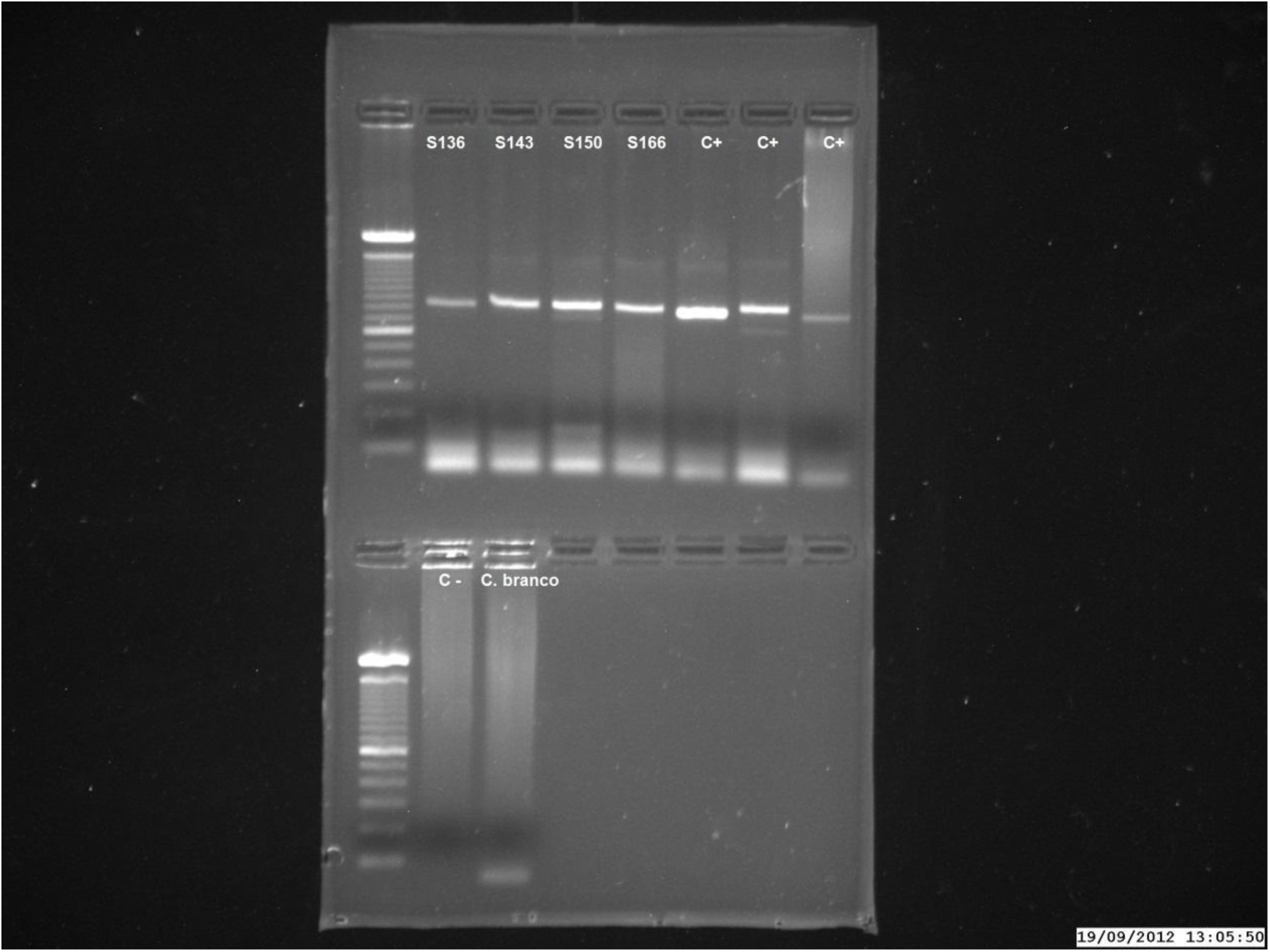
An example of a PCR Gel of samples S136 (*Sapajus apella*), S143 (*Sapajus apella*), S150 (*Pithecia irrorata*) and S166 (*Alouatta puruensis*) that tested positive for HBV and whose products were then sequenced.

Sequence analysis of a 547-base pair (bp) fragment of the S gene in these five positive samples showed that four sequences (from *Alouatta puruensis, Ateles chamek, Pithecia irrorata,* and *Sapajus apella*) belonged to the human HBV-A genotype clade and one (from *Callicebus bruneus*) to the human HBV-D clade (Fig 3), which are all distinct from the HBV genotypes isolated from apes (Fig 4), and they shared very high sequence similarity with human HBV sequences.

**Fig 3.**
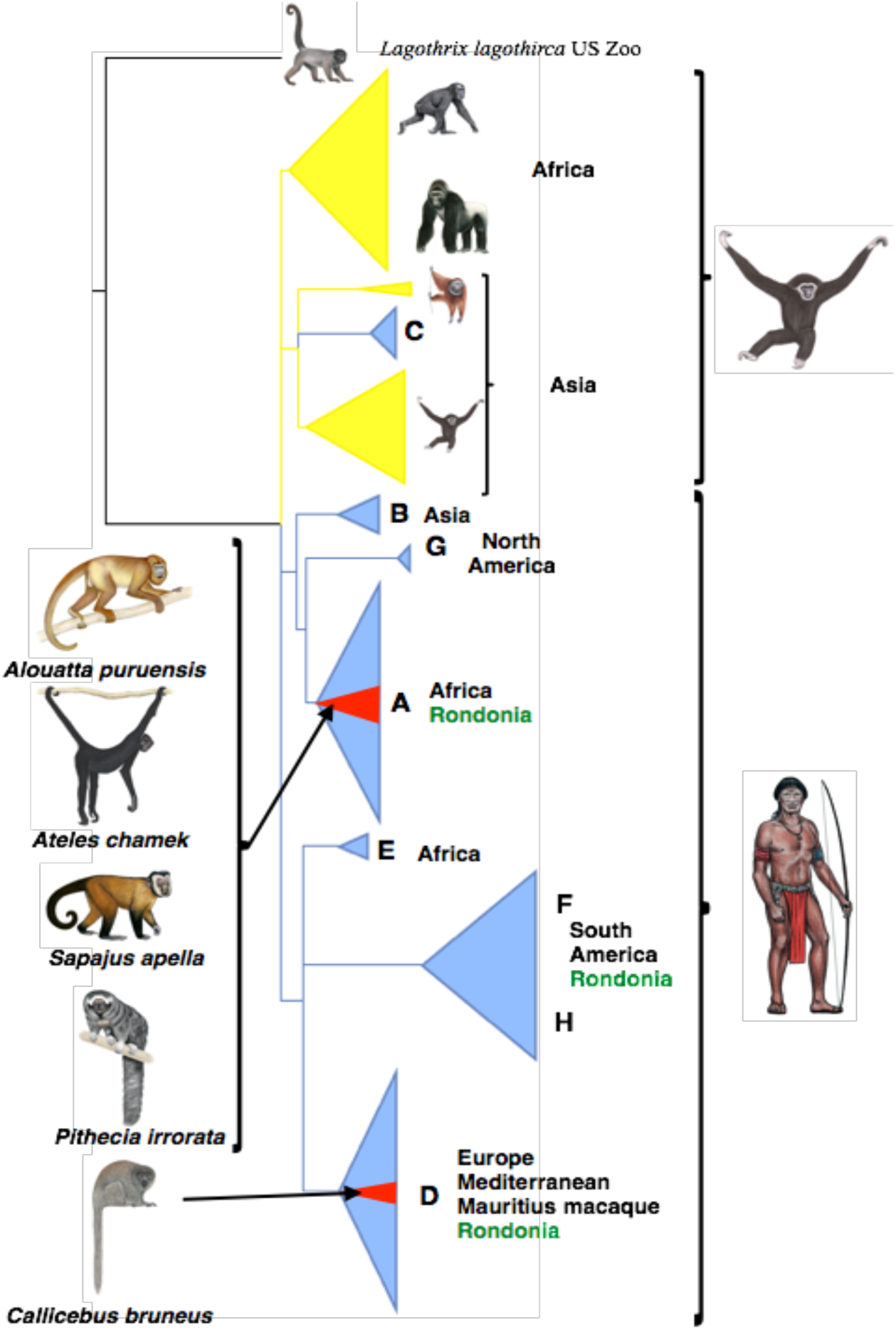
Bayesian tree based on a 547-bp fragment of the S gene for HBV different strains from humans (in blue), and African and Asian apes (in yellow). Our five New World primate samples from Rondônia are shown as red triangles within the collapsed clades. The Human HBV types found in Rondônia are shown in green. Based on Fig 4. Primate Illustrations copyright 2014 Stephen D. Nash/Conservation International/ IUCN SSC Primate Specialist group. Used with permission.

**Fig 4.**
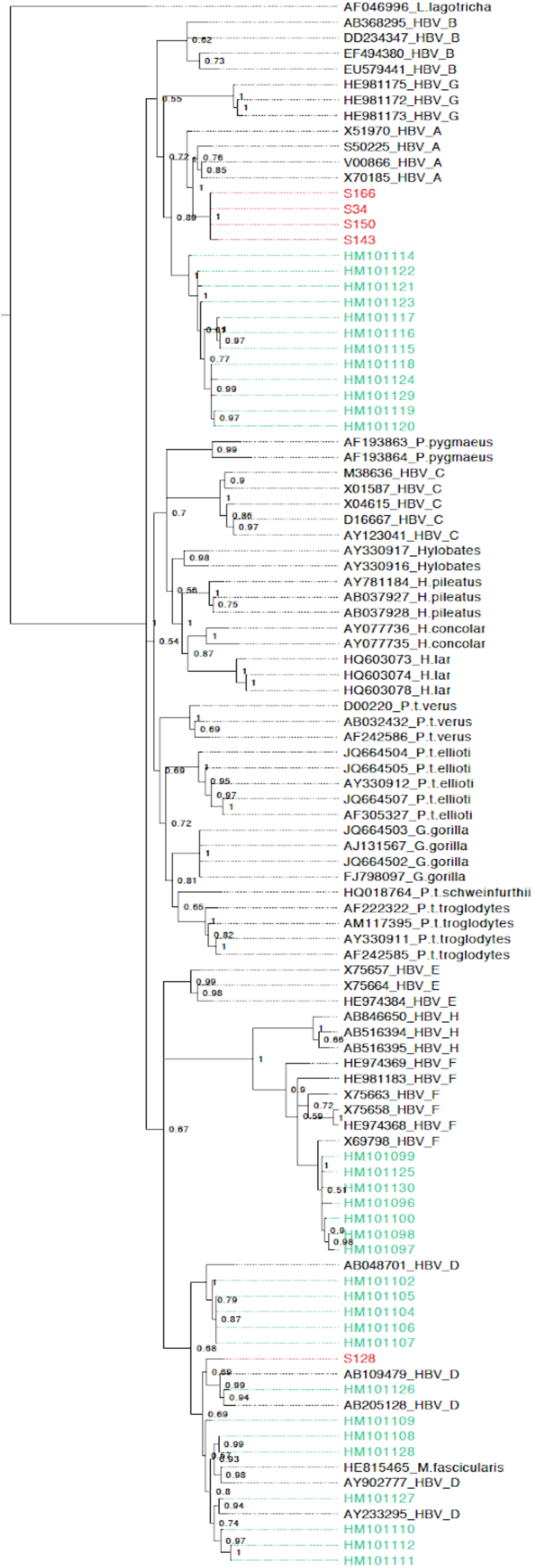
Bayesian tree based on a 547-bp fragment of the HBV S gene of different strains of the virus from Humans alongside Apes from African and Asian. Our five New World Primate samples from Rondônia are shown in red and sequences from Human Rondônia patients are shown in green. Node values correspond to the probability.

## Discussion

Colonization and immigration in southwestern Amazonia (in Rondônia state) began in the 17th century, driven by different development cycles: mining, slavery, rubber production, railway development, hydroelectric dams. More recently, the region also received immigration of international refugees. These migration waves created a multicultural population [19], where 57.7% are non-native inhabitants [20]. In that region, the HBV-A genotype was the most prevalent strain in humans followed by the D-strain [18, 21, 22]. This reflects the movement of tens of thousands of workers of African descent into this area, initially to help construct the Madeira-Mamore railway between 1907 and 1912, before the arrival of hundreds of thousands of Europeans to farm the land. Those strains match with those found in our primate samples, which support the hypothesis of a human-primate transmission of HBV. To our knowledge, our study represents the first reported cases of HBV in multiple wild New World primate species anywhere in the world.

The possibility of human-primate spillover is also supported by the analysis of 39 liver tissues from 11 species monkeys collected from a very remote and pristine region of the Amazon that is mostly unaffected by humans, i.e., the upper Japura River. None of these monkeys tested positive for HBV, thereby suggesting that humans are the source of HBV in New World primates. According to Dupinay et al. [13], the HBV strain found in long tail macaques in Mauritius also originated from a human source. Thus, although apes have had time to develop specific strains of HBV, evidence indicates that only human strains of HBV have been found in both Old World and New World monkeys, which strongly suggests that HBV was transmitted to them by humans. HBV has been detected in the saliva of HBV carriers [23, 24] and it is a proven transmission route between humans [25] and from humans to gibbons under laboratory conditions [26, 27]. However, the present study provides the first evidence for the natural human to primate transmission of HBV across multiple genera, thereby supporting the idea of human to ape transfer in the Old World [5]. The HBV strain isolated from a woolly monkey at Louisville Zoo was not related to any known HBV strain. Hu et al. [28] reported that this strain was related to the human HBV-F strain, but our analysis supports an earlier study [15], which suggested that this is the most differentiated and basal form among primate HBV strains. In the present study, both of our woolly monkey samples tested negative.

In tropical countries, high levels of forest clearance and human settlement to allow large-scale cattle ranching, other types of agriculture, and infra-structure projects are bringing humans into direct contact with wildlife. Areas of interface between primary forest and human habitation allow very close physical contact between humans and nonhuman primates. Our results suggest that the impact of the human occupation of the Amazon extends far beyond habitat and biodiversity losses. Human population density was found to be an important driver of primate HBV infections, which reflects that human occupation has been increasing the interactions between human and wildlife and facilitating the cross-transmission of diseases. In particular, diseases such as HBV and human malaria have been introduced by human settlers and they may pose major threats to local wildlife. Indeed, monkeys are important targets of hunters in the Amazon, and baby monkeys are commonly kept as family pets, thereby increasing the possible cross-species transmission of viruses such as HBV that is known to share hosts in nature and domesticated species [29]. For instance, domesticated fowl and swine in China were both found to be naturally infected with HBV [29], while swine in Brazil were infected with HBV that shared 93%-96% sequence identity to human HBV [30].

Of greater concern is the possibility that a more virulent HBV strain might spill back into the human population [31], especially in areas with increasing human occupation such as the Amazon, Asia, and Africa. The transmission of HBV from primates to humans can occur through wild meat hunting and trade. When people handle, butcher, consume or trade primate meat, they may be exposed to infected bodily fluids, such as blood, which is a known vector for HBV transmission. This risk is exacerbated in areas like the Amazon, where primates are a significant target of these activities. For instance, it is estimated that c. 1,500 kg of primate meat are consumed annually in five cities in central Amazonia [32]. In addition, primates such as squirrel monkeys (*Saimiri* spp.) are commonly traded as pets and considered to be among the most profitable wildlife species sold in wildlife markets in Amazonia [33].

Rondônia is one of Brazil’s nine Amazonian states and dramatic changes in the natural cover have occurred in this area over the past 30 years, where the forest losses have been estimated at up to 80% in some municipalities [34, 35]. Experimental observations have shown that chimpanzees infected with human HBV infection may develop clinical symptoms, such as anorexia, lethargy, and jaundice. Furthermore, the primate species that tested positive for HBV in the present study are reservoirs of human malaria in Rondônia due to their high rate of infection with *Plasmodium* and their proximity to urban areas [36].

## Materials and Methods

Primates were sampled in two separate sites of the Brazilian Amazon between July 2009 and November 2013: 1) throughout the southwestern Amazonia (i.e., municipalities in the state of Rondônia and two nearby cities in Mato Grosso and Amazonas state), a region with high human occupation (i.e., 1,777 million inhabitants), urbanization, and deforestation (i.e., around 31% of its native vegetation lost in the last 30 years); and 2) the upper Japura River region, a remote area deep in the Amazon that is highly pristine (i.e., mostly unaffected by humans). Animals were all wild and either captured live to obtain blood samples (Rondônia: 49 samples of 17 species) or collected as voucher specimens to acquire liver tissues (Japura: 39 samples of 11 species) (Table S1) and represented a wide range of species (Fig 5 A).

**Fig 5.**
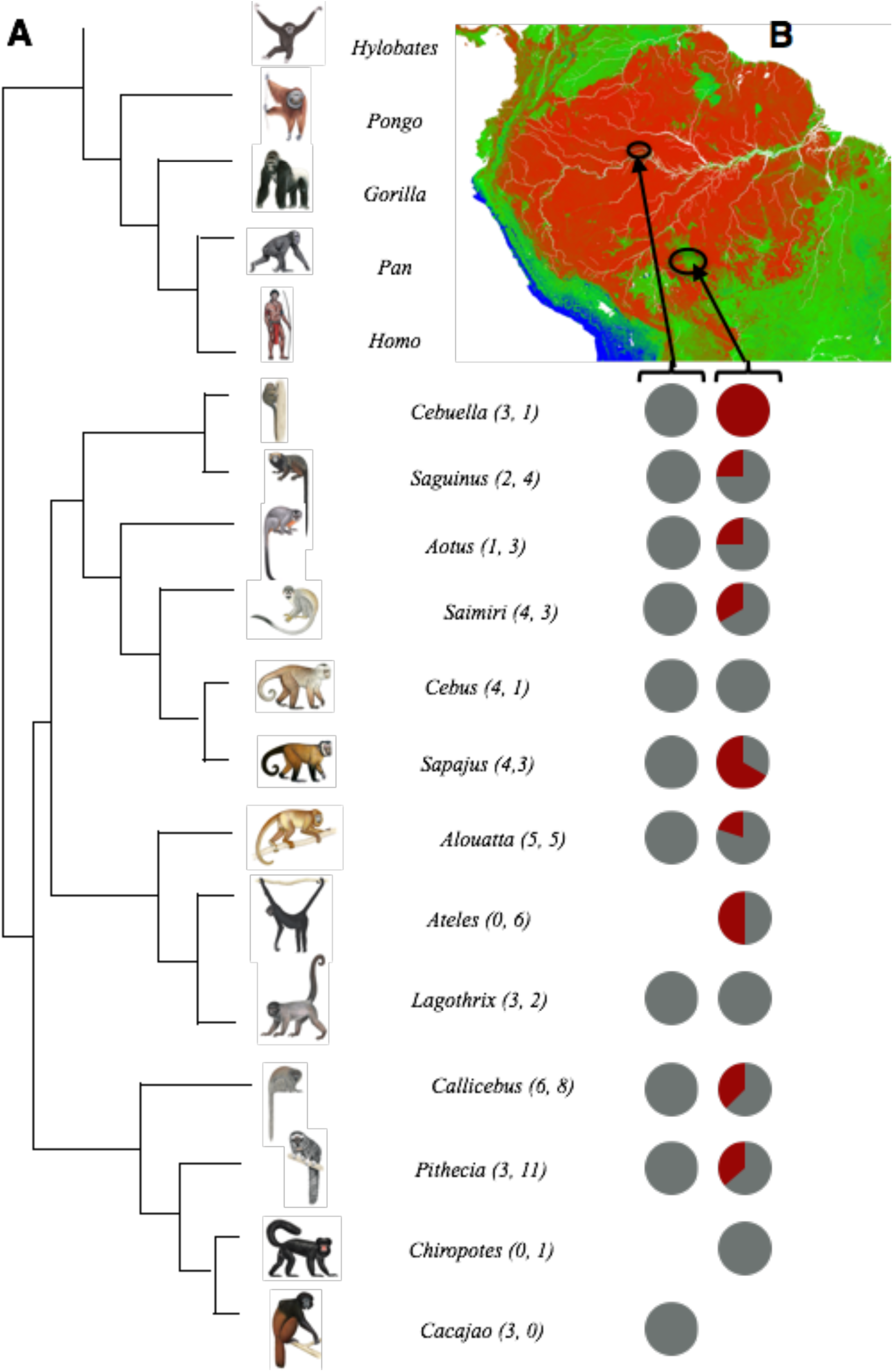
**A.** Phylogenetic relationships among all the primate genera considered in this study based on that of Perelman *et al*. [37]. **B.** Map of the northern region of South America showing the locations where the samples were collected. Red represents dense Amazonian forest and green denotes open areas, including areas of anthropic deforestation (source: NASA Observatory). The pie charts show the percentages of sampled primates that tested positive for HBV in the Japura River area (left) and Rondônia (right). Grey denotes negative PCR reactions and red indicates positive PCR reactions. Each group of pie charts represents individuals in the corresponding primate genus shown on the left in part **A**.

Details of the sampling procedures and protocols are described elsewhere [36]. We extracted viral DNA from 200 μL of blood serum using an *in-house* method based on membrane lysis with guanidine isothiocyanate and chloroform, which was adapted from a previously reported method [38]. DNA was extracted from all liver tissues using a standard phenol-chloroform protocol [39]. Following DNA extraction, amplification was performed by duplex polymerase chain reaction (PCR) to generate amplicons of 1.5 kilobases (kb) and 1.7 kb, as described by Tran *et al*. [40]. If these two bands were not detected by agarose gel electrophoresis, the amplicons were subjected to nested PCR amplification of a 741-bp fragment of the rt/S gene using RT/S forward (5′-AAGGTATGTTGCCCCGTTTGTC-3′) and RT/S reverse (5′-GGGTTGCGTCAGCAAACACT-3′) primers, according to Gauthier *et al*. [41]. All the PCR products were separated by agarose gel electrophoresis. Positive PCR products were purified using an ExoSAP-IT kit (GE, USA) according to the manufacturer’s instructions, and then sequenced with an ABI Prism® 377 Automatic Sequencer (Applied Biosystems, Foster City, CA, USA).

All the sequences were confirmed by eye using ProSeq 3.0 [42]. A 547-bp fragment of the S gene was used in the final phylogenetic analysis, which included three additional datasets. The first comprised representative sequences for human HBV genotypes A–H (**A:** V00866, X51970, X70185, S50225; **B:** U95551, X97848, X85254, X65257; **C:** D16665, D50489, D16666-7, D50517, D12980, D50519, AY123041, D50520, D00630, D23682, X52939, V00867, X04615, M38636, D23684, X01587, D28880, D23680-1; **D:** D00329-31, M54923, X97850-1, D23678-9, D50521-2; **E:** X75664, HE974384, X75657; **F:** HE981183, X75663, X75658, HE974368-9, X69798, **G:** HE981172-3, HE981175; **H:** AB516394-5, AB846650). The second comprised HBV sequences from gorillas (AJ131567, FJ798095-7, JQ664502-3), chimpanzees (D00220, AB032432-3, AF222322-3, AF242585-6, AF305327, AF498266, AM117395-7, AY330911-2, FJ798098-9, HQ018763-4, JQ664504-9), gibbons (AB037927-8, AY077735-6, AY330915-7, AY781184), orangutans (AF193863-4) and the woolly monkey (AF046996) from Louisville Zoo.

We employed a generalized linear mixed model (GLMM) to investigate the influence of human population density (persons/km²), deforestation level (percentage of native cover loss), and Gross Domestic Product per capita (GDP per capita, in US$) on the likelihood of primates testing positive for HBV at different sampling sites. We treated HBV status as a binary response variable, coding positive samples as 0 and negative samples as 1, stratified by species and sampling site. The model was run using the *gamlss* package in R studio (Version 2023.09.1+494) with a Binomial distribution family. Our approach involved executing a stepwise regression procedure, incorporating both backward and forward selection methods. This stepwise process iteratively adds or removes explanatory variables based on their contribution to enhancing the model’s fit. We incorporated primate species as a random factor to account for unmeasured species-specific effects, such as varying susceptibilities to HBV influenced by genetic, physiological, and behavioral factors. This provided a more accurate and generalizable understanding of the factors influencing HBV prevalence across diverse primate species. In addition, we also applied the ‘weights’ term to adjust for disparities in sample sizes across different primate species and collection sites, ensuring a more balanced and representative model outcome.

The selection of variables was done by the Akaike Information Criterion (AIC). The AIC not only considers the goodness of fit but also incorporates a penalty for the number of parameters in the model, thereby balancing model complexity against the risk of overfitting. In each step of the regression, the AIC of the model was calculated, and the variable whose addition or removal resulted in the lowest AIC was chosen for the next iteration. This process continued until no further reduction in AIC was possible, at which point we obtained our final model.

Finally, human strains and genotypes from patients in Rondônia (HM101096-100, HM101102, HM101104-12, HM101114-30) were compared directly with our primate samples to determine the genotypes of HBV strains infecting primates. Sequences were aligned using CLUSTALX 2.1 [43]. The most appropriate evolutionary model for this dataset was defined as GTR+I+G based on Akaike’s information criterion [44] using MrModeltest 2.3 [45]. Bayesian analysis was performed using MrBayes 3.2 [46] for 10,000,000 generations with sampling every 100 generations, where the first 10% were discarded as the burn-in.

Strict procedures were followed to avoid false positives [47], including splitting the samples and performing extraction procedures at two different times. All procedures adopted in this study were approved, in full compliance with specific federal guidelines issued by the Brazilian Ministry of Environment (SISBIO, process number 14081–3 and 17302–1).

## Acknowledgments

We thank Santo Antônio Energia S/A and Energia Sustentável do Brasil (ESBR) for permission to capture animals in their reserves. We thank Fabricio Bertuol for carrying out some of the molecular analyses in Manaus. Molecular analyses and Japura field expeditions were funded by CNPq SISBIOTA Program (No. 563348/2010-0) to IPF, and in Rondônia by grants from the Fundação Oswaldo Cruz Rondônia (FIOCRUZ/RO), Fiocruz-Rondônia, CT-Amazônia, and CNPq Universal no. 471196/2011-8. Permission to conduct fieldwork and to collect tissue samples was granted by IBAMA and ICMBio (License No. 005/2005 – CGFAU/LIC).

## Supplemental data

**Table S1:**
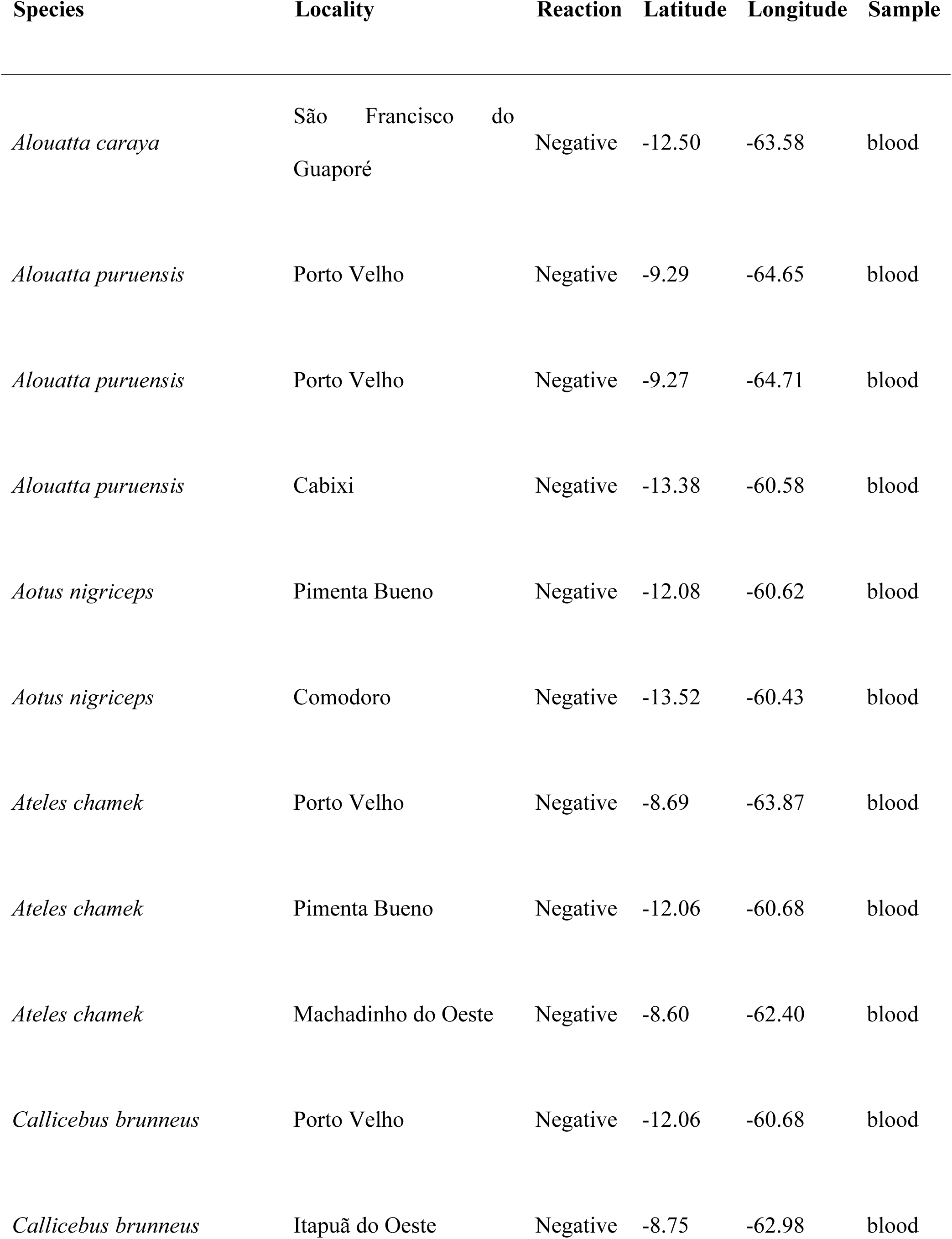
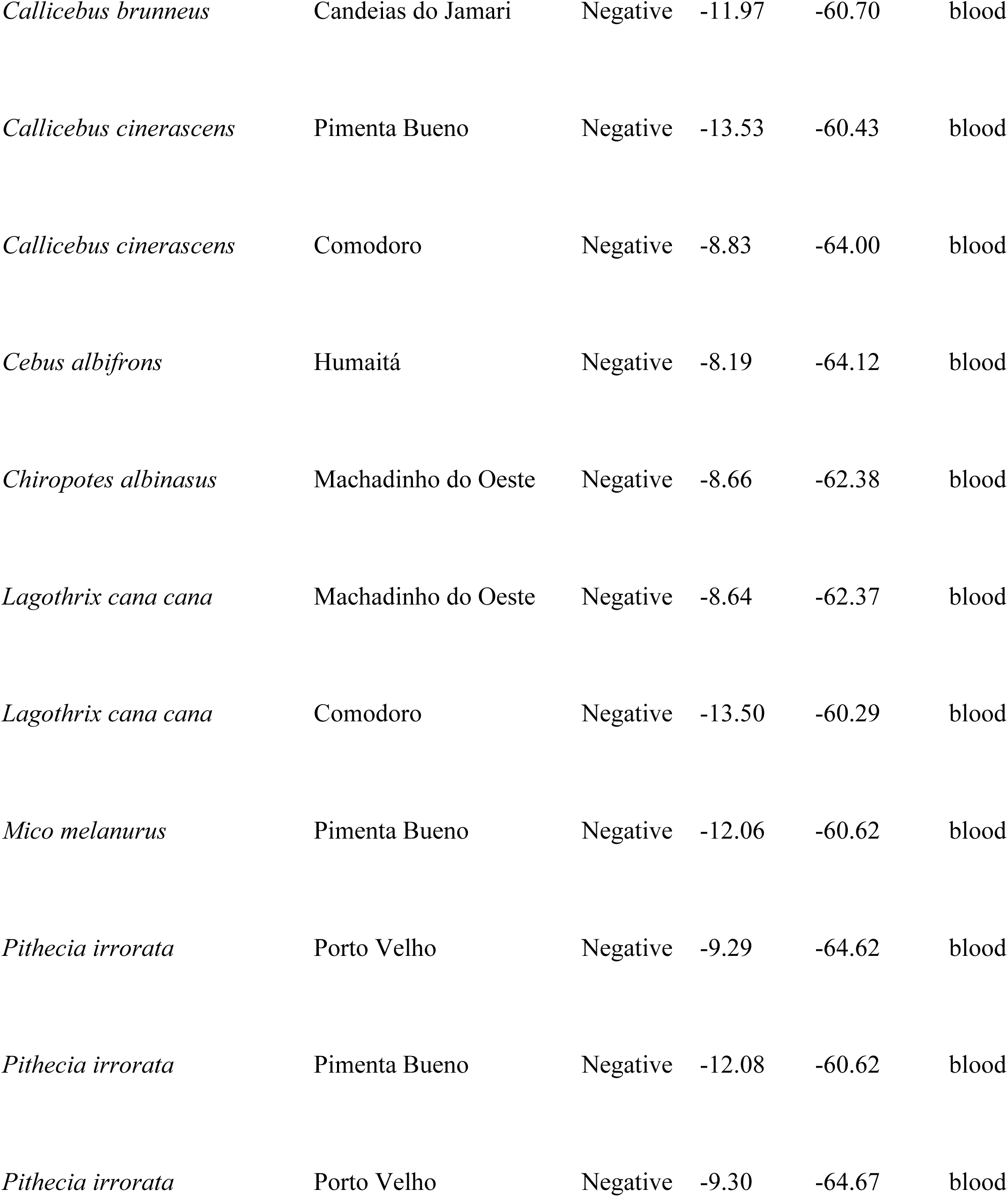
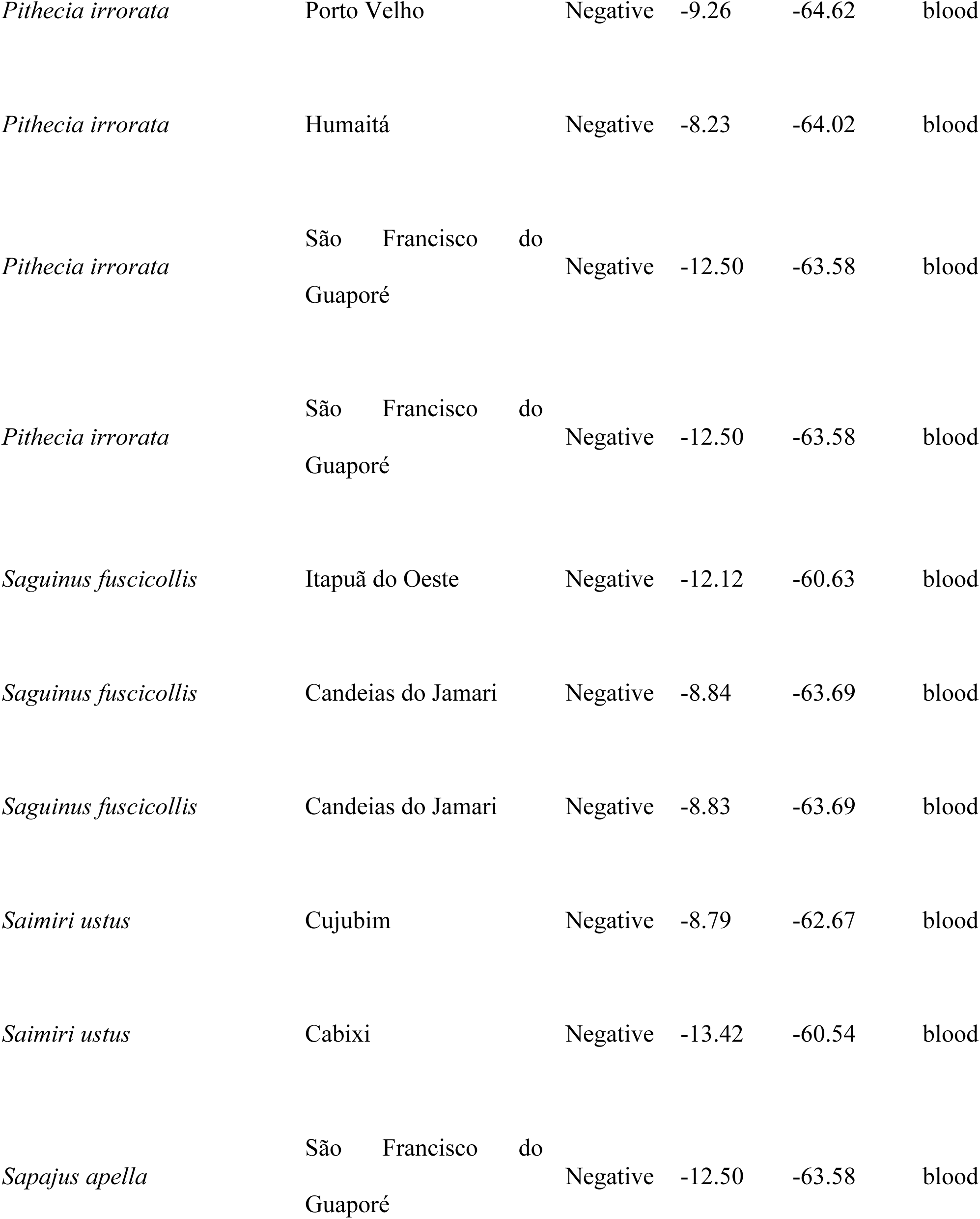
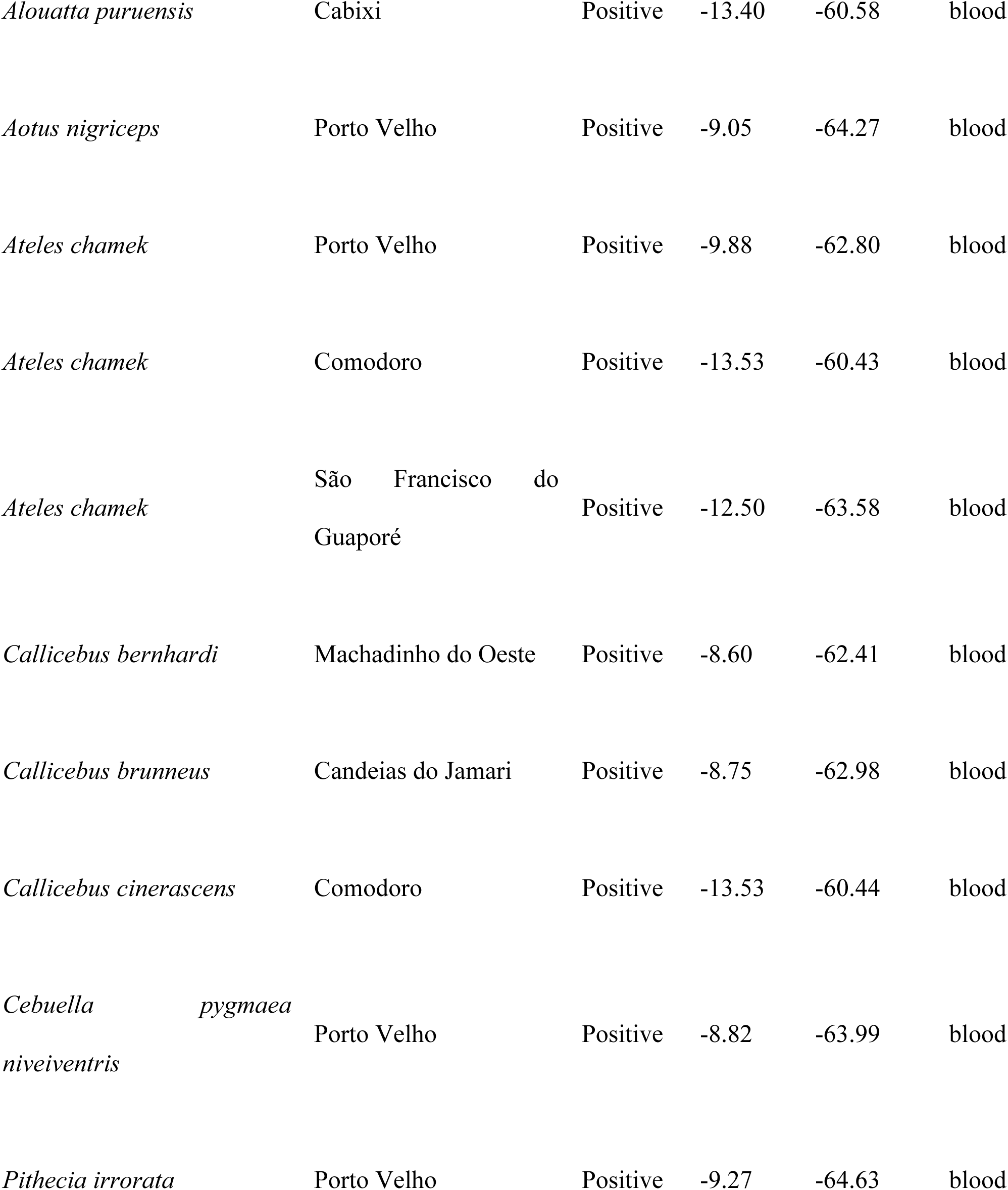
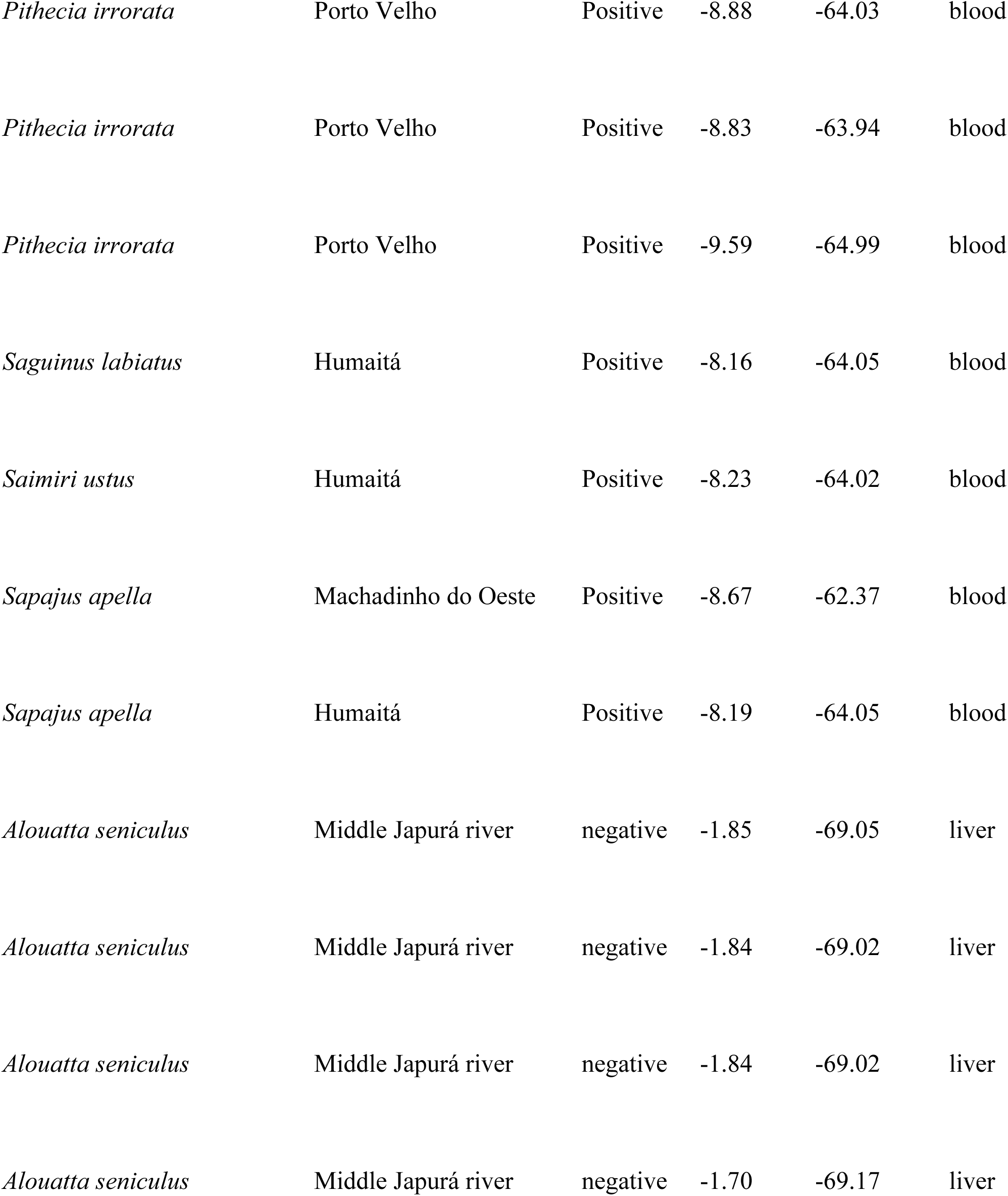
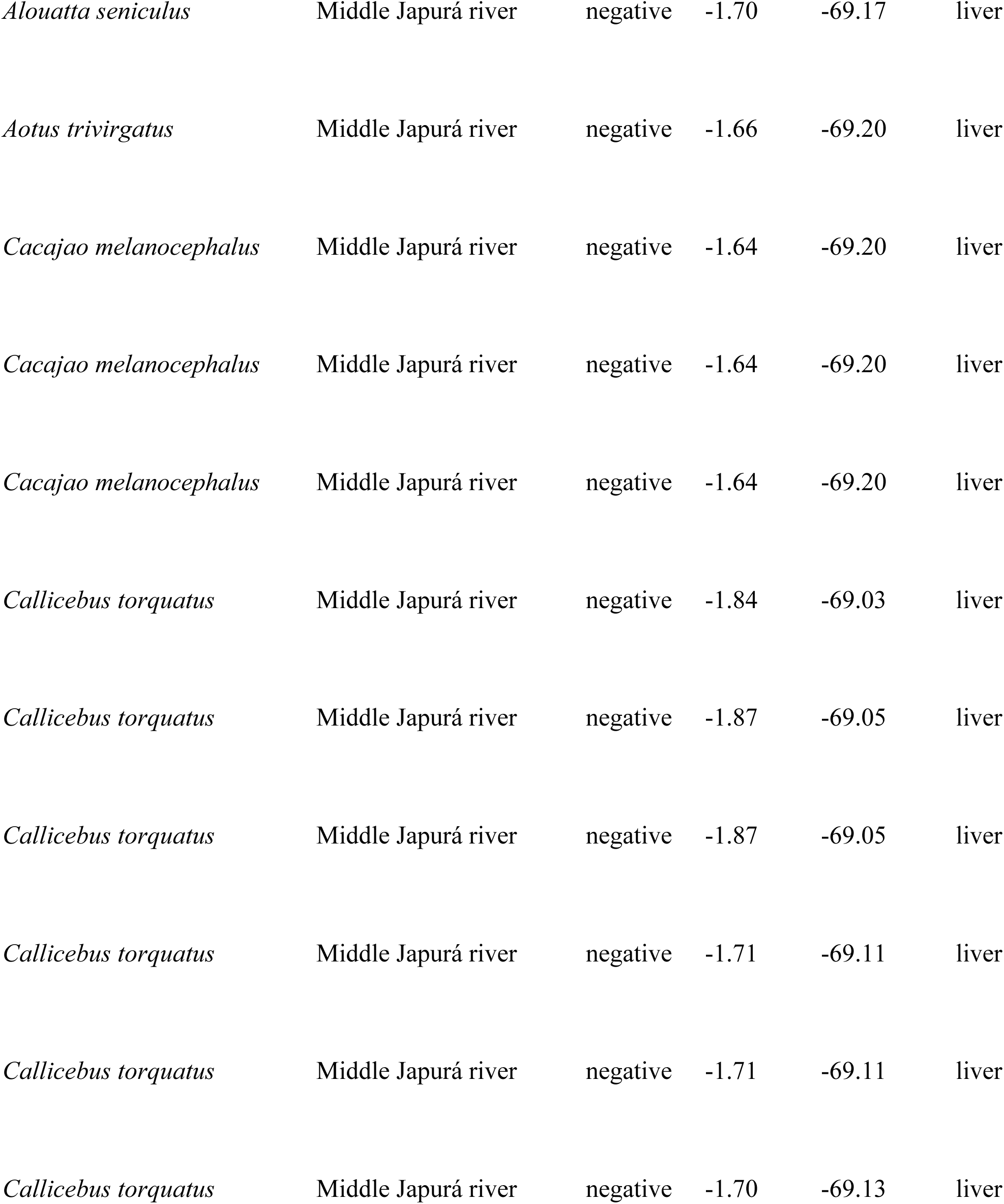
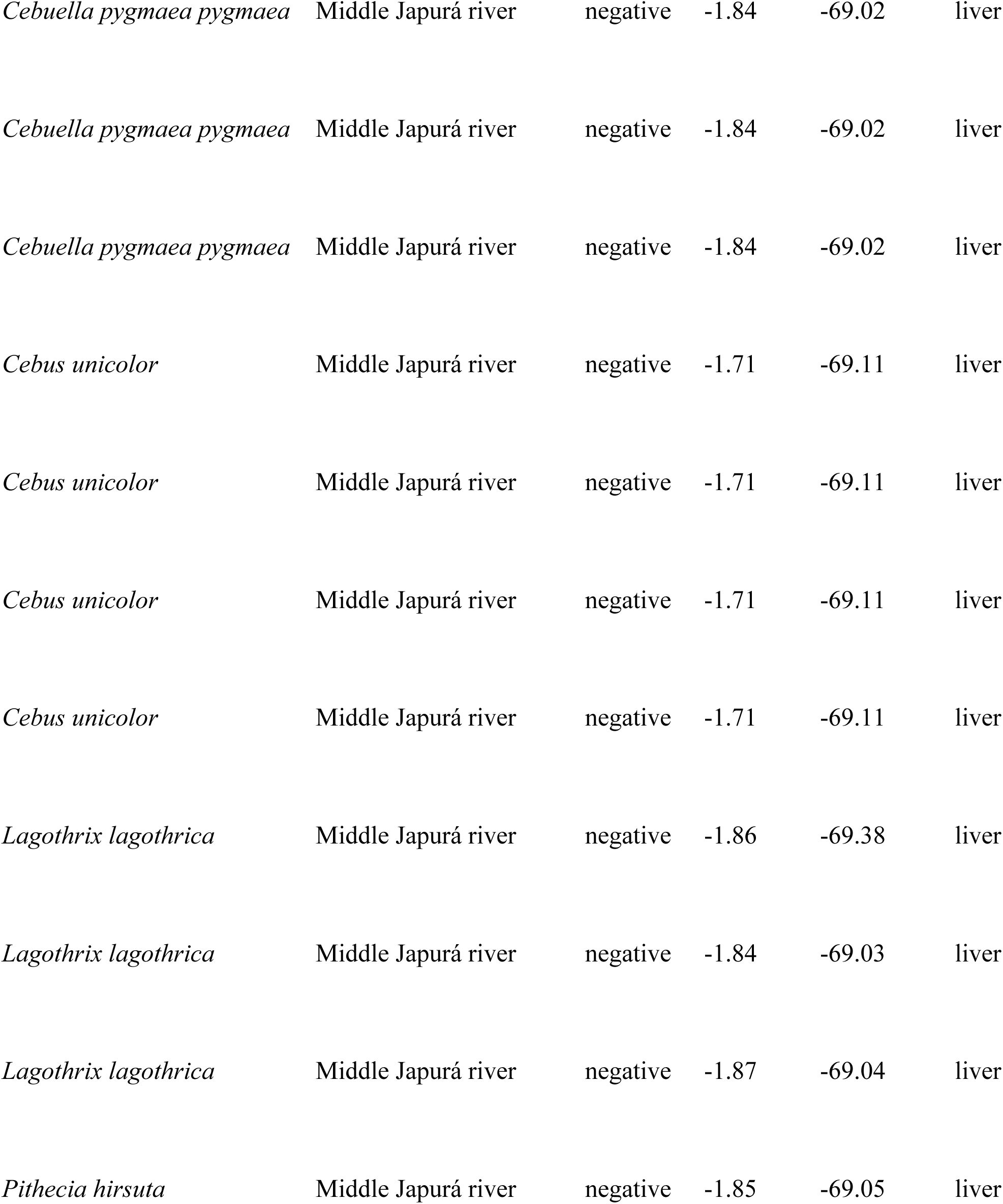
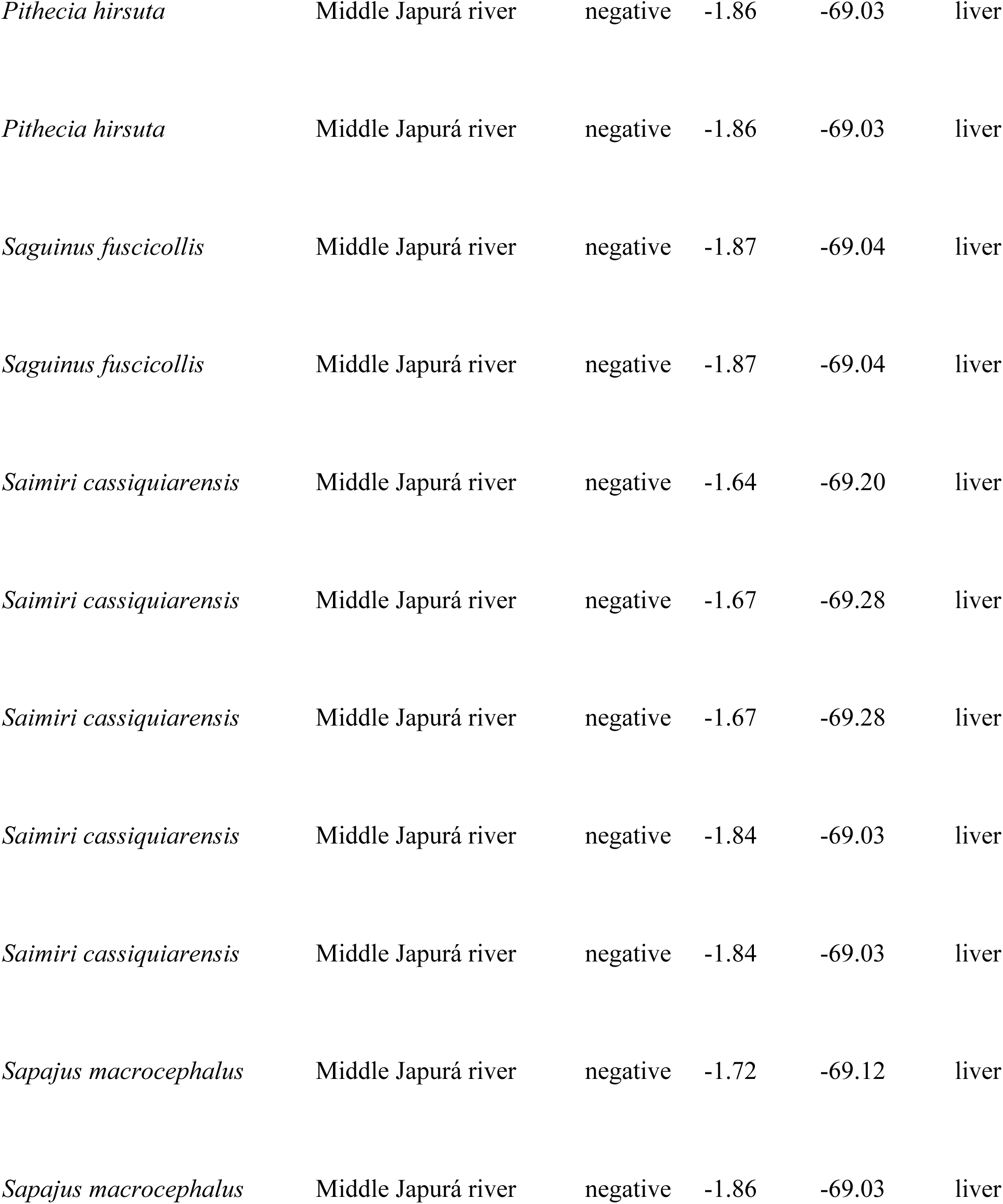
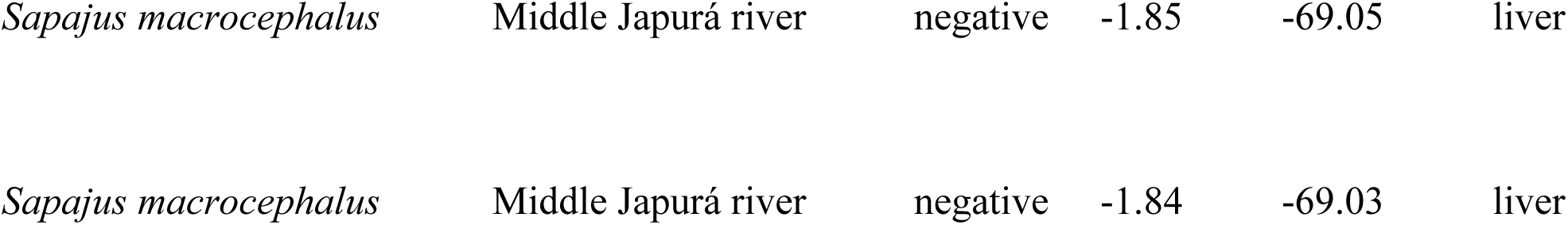
List of wild New World Primate samples from specimens caught in Rondônia and Japurá areas, Brazil; result of hepatitis B (reaction) in two different types of samples.

## Relevant data underlying the research

In this link we include the full sequence alignment used in this study: https://drive.google.com/file/d/1VwrY9Q1JXVCDCm2MYpfnUUQE8c5pYmYC/view?usp=drive_link

